# Structure-based virtual screening and Molecular Dynamic Simulations identified FDA-approved molecules as potential inhibitors against the surface proteins of H1N1

**DOI:** 10.1101/2023.12.02.569695

**Authors:** Sukanya Bordoloi, Roshny Prasad, VS Lakshmi, Rajesh Chandramohanadas, Kathiresan Natarajan, Shijulal Nelson-Sathi

## Abstract

H1N1, a subtype of influenza A virus, remains a global concern due to its ability to cause highly contagious, and fatal infections of the respiratory tract that can lead to seasonal pandemics. Factors such as genetic reassortment and antigenic shift due to the segmented nature of its RNA genome, lead to its rapid evolution that impacts drug and vaccine interactions. Thus, there is growing interest in identifying small-molecules against H1N1 to address the challenges posed by its ever-changing nature and enhance ability to combat the virus. By targeting the surface proteins crucial in the viral life cycle and interactions with host cells, we conducted structure-based virtual screening of 2,471 FDA-approved small molecules against H1N1’s hemagglutinin (HA) and neuraminidase (NA) using molecular docking, and molecular dynamic simulations. A binding pocket for HA was identified at the interface of the three protomers, close to the fusion peptide, while in the case of NA the co-crystal ligand binding site was targeted. We identified 5 molecules, namely Econazole, Butoconazole, Miconazole, Isoconazole, and Tioconazole with higher binding affinity to HA and 4 molecules, namely Acarbose, Rutin, Paromomycin, and Idarubicin showing superior binding affinity to NA. Further molecular dynamic simulation of these molecules bound with HA and NA reaffirm the stability of the complexes. These molecules are known to have antifungal and antiviral properties. Thus, this study elucidates the importance of targeting HA and NA and paves the way for repurposing existing antivirals, antibacterials, and antifungals as inhibitors of the H1N1 viral entry into host cells.

## Introduction

The emergence of the novel pandemic-causing influenza A (H1N1) virus in Mexico in April 2009 turned out to be a significant global health event (Cox & Subbarao, 2000; Gatherer, 2009; Mena et al., 2016). It rapidly spread across the countries, resulting in an estimated 201,200 respiratory disease-related deaths worldwide, and continues to circulate as a major constituent of seasonal influenza viruses (Dawood et al., 2012; Jones et al., 2019; WHO, 2023). The H1N1 viral genome consists of eight negative-sense single-stranded RNA segments encoding 12 viral proteins, including the envelope glycoproteins hemagglutinin (HA) and neuraminidase (NA) (Dadonaite et al., 2019). Viral entry mediated by HA takes place through two major steps (Danieli et al., 1996). The first step is the recognition of the target host cell by binding of HA to sialic acid-containing receptors on the host cell surface which causes the virus particle to be internalized via endocytosis. The next step is the fusion of the viral and endosomal membranes which is brought about by the pH-induced conformational change of the HA trimer. The key player during this process is the fusion peptide, a short hydrophobic region of about 23 amino acid residues. It undergoes a loop-to-helix transformation that enables it to interact with the lipid bilayer of the endosomal membrane. The role of NA is to complete the infectious cycle. It is a sialidase that cleaves the alpha-ketosidic bond between sialic acid and the adjacent sugar residue in order to prevent the aggregation of newly formed virions and re-binding of the virion to the infected cell via HA (Shtyrya et al., 2009). The abundance of HA and NA on the viral surface makes them suitable drug targets in the current scenario (Bai et al., 2021).

Antiviral therapy is a widely accepted intervention method against respiratory disease-causing viruses like H1N1 and most of the antivirals approved by the Food and Drug Administration (FDA) are small molecules (Naesens et al., 2016; Chaudhuri et al., 2018). Currently, M2 ion channel inhibitors (amantadine, rimantadine), and neuraminidase inhibitors (zanamivir, oseltamivir, peramivir) are approved by the FDA against influenza infections (Batool et al., 2023). However, the emergence of drug-resistant variants has rendered the M2 ion channel inhibitors obsolete (Hurt et al., 2012). There have also been reports of viral resistance against neuraminidase inhibitors (Collins et al., 2009; Mckimm-Breschkin, 2013). At present, there are no FDA-approved drugs against HA, however it is of great interest to researchers owing to its role in viral entry. Apart from certain sialic acid analogs and pentacyclic triterpenoids, the discovery of small molecule inhibitors of the HA receptor binding pocket has been challenging so far (Wang et al., 2013; Yu et al., 2014). However, several small molecules have been reported to bind in the HA stem region, inhibiting the fusion step (Chen et al., 2021).

The stem region of HA, which mainly comprises of the HA2 subunit can be a suitable target to exploit due to its conserved nature in comparison to the head region made of the HA1 subunit (Fan et al., 2015). In our study, we have targeted this stem region of the functional pre-fusion homotrimeric form of HA and have done a structure-based in-silico screening of FDA-approved drug molecules against a hydrophobic pocket which is in the close vicinity of fusion peptides of all three protomers. We have also screened these molecules against the substrate binding site of NA near the 150-loop region. Such a drug repurposing approach can be a time-efficient way to meet the increasing demand for H1N1 inhibitors.

## Materials and Methods

### Target Selection and Protein Preparation

The crystal structures corresponding to A/California/04/2009 H1N1 hemagglutinin (wild-type HA trimer) and neuraminidase (NA) were retrieved from Protein Data Bank (PDB) (HA-3LZG by Xu et al., 2010; NA-3TI6 by Vavricka et al., 2011). In this study, the HA protein was considered in its functional trimeric form to identify the optimal site for targeting, crucial for inhibiting membrane fusion. Conversely, for the NA protein, a monomeric structure complexed with oseltamivir carboxylate (the co-crystallized ligand) was selected for further analysis. All the non-protein entities in the initial HA and NA crystal structures were meticulously removed, with the exception of the co-crystallized ligand and the calcium ion situated proximal to the active site in NA, as previously reported (Chong et al., 1991). Prior to the molecular docking procedures, both protein structures underwent comprehensive preparation using the Protein Preparation Workflow module within Schrödinger Maestro Version 13.5 (Sastry et al., 2013). This preparatory phase encompassed the addition of missing hydrogen atoms, elimination of co-crystallized water molecules, assignment of partial charges, and application of restrained minimization employing the OPLS4 force field after the pre-processing to ensure the optimal geometrical and energetic state of the proteins (Lu et al., 2021).

### Binding Site selection and receptor grid generation

To identify potential binding sites on the HA protein, the SiteMap tool of Schrödinger Maestro Version 13.5 was employed (T. Halgren, 2007; 2009). This approach facilitated the prediction of 5 druggable pockets, with each site map delineated within a 4 Å radius from the nearest site point. The pocket exhibiting the highest SiteScore was subsequently chosen for further investigation. The corresponding grid file was generated using the Receptor Grid Generation panel in the Glide module of Schrödinger Maestro Version 13.5, employing default parameters, in order to define the three-dimensional space around the protein where the ligands will be positioned during the docking calculations. This grid file was then utilized in the ensuing molecular docking protocol. In the case of the NA protein, the binding site associated with the co-crystal ligand was directly selected. The corresponding grid file was generated using default parameters for subsequent molecular docking studies.

### Ligand preparation

A set of 2,471 FDA-approved small molecules were downloaded from the DrugBank database in SMILE format and ligand preparation was performed using the LigPrep module in Schrödinger Maestro version 13.5 (Wishart et al., 2006; LigPrep, Schrödinger, LLC, New York, NY, 2017. New York, NY.). Through this step, the tautomeric and ionization states of the compounds were expanded to reflect the potential variations that may occur under physiological conditions, ring conformations and stereoisomers were produced to account for the flexibility and potential binding modes of the ligands.

### Molecular docking and MM-GBSA calculations

To evaluate the ligand library against the identified binding sites in both HA and NA proteins, a three-tiered rigid receptor docking was carried out. This involved consecutive runs in high-throughput virtual screening (HTVS), standard precision (SP), and extra precision (XP) modes, using the Glide module within Schrödinger Maestro 13.5 (Friesner et al., 2004, 2006; T. A. Halgren et al., 2004). Following the docking runs, molecular mechanics-generalized Born surface area (MM-GBSA) calculations were performed for the rescoring of the docked ligand poses. This step enabled the computation of binding free energies and the estimation of relative binding affinities (Guimarães &amp; Cardozo, 2008). The Prime module within Schrödinger Maestro 13.5 was utilized for this purpose, using the VSGB 2.1 implicit solvation model and the OPLS4 force field with default parameters (J. Li et al., 2011). The selection criteria for further assessment were set at docking scores below −7 and binding free energy values lower than −40 kcal/mol. This threshold ensured that only the most promising ligand candidates were considered for subsequent analyses. Additionally, to validate the docking protocol and confirm its accuracy, the co-crystal ligand (oseltamivir carboxylate) bound to NA (PDB ID: 3TI6) was re-docked using the same protocol. Finally, the resulting docked complex structures were visualized and analyzed using Pymol, providing a detailed overview of the ligand-protein interactions (Delano, 2002).

### Molecular dynamics simulation

MD simulations were conducted over a 100 ns time scale utilizing the Desmond module in Schrödinger Maestro 13.5 (Bowers et al., 2006). Prior to the simulations, a simulation box was constructed for the chosen protein-ligand complexes using the System builder tool in Desmond. The complexes were immersed in a TIP3P water model within an orthorhombic box, ensuring a 10 Å extension beyond the outermost atoms of the complex for accurate representation (Mark &amp; Nilsson, 2001). To maintain system neutrality, an appropriate number of counter ions were added. The MD simulation was carried out in the NPT ensemble at a constant temperature of 300 K and a pressure of 1.63 bar, over the 100 ns duration. The OPLS4 force field was employed to ensure a reliable representation of the intermolecular forces and energies during the simulation (Lu et al., 2021). For visualization and data analysis, the Simulation interaction diagram tool within Maestro was used to generate plots and figures. In addition to MD simulations, Principal Component Analysis (PCA) was carried out to extract essential dynamic information from the generated trajectories by reducing the dimensionality of simulation data and revealing the dominant modes of motion within the system (David & Jacobs, 2014). This was executed using the trj_essential_dynamics.py script in Schrödinger’s command-line mode. Free energy landscapes (FEL) of each simulation were calculated using the Geo-measures plugin (Kagami et al., 2020). The MD trajectories against RMSD and Radius of Gyration (RG) were plotted as 2D graphs. Geo-measures consists of the powerful module g_shad of the GROMACS program.

## Results and discussion

### Ligand Binding Site of HA

In order to perform molecular docking, a hydrophobic pocket at the interface of the stem region of the three protomers of HA was identified. This pocket is made up of amino acid residues of the fusion domain (V-29 and F-294 of the HA1 subunit), fusion peptide (L-2, F-3 of HA2 subunit), and HA2 ectodomain (K-51, W-92, Y-94, N-95, A-96, L-98, L-99, L-102, E-103, E-105, R-106, T-107, D-109, and Y-110) (Figure 1). The pocket had a site score of 1.22 and druggability score of 0.96. It also showed less solvent exposure (exposure score= 0.31, enclosure score=0.96) and had a considerable size (size= 216, volume= 335.11).

**Figure 1:**
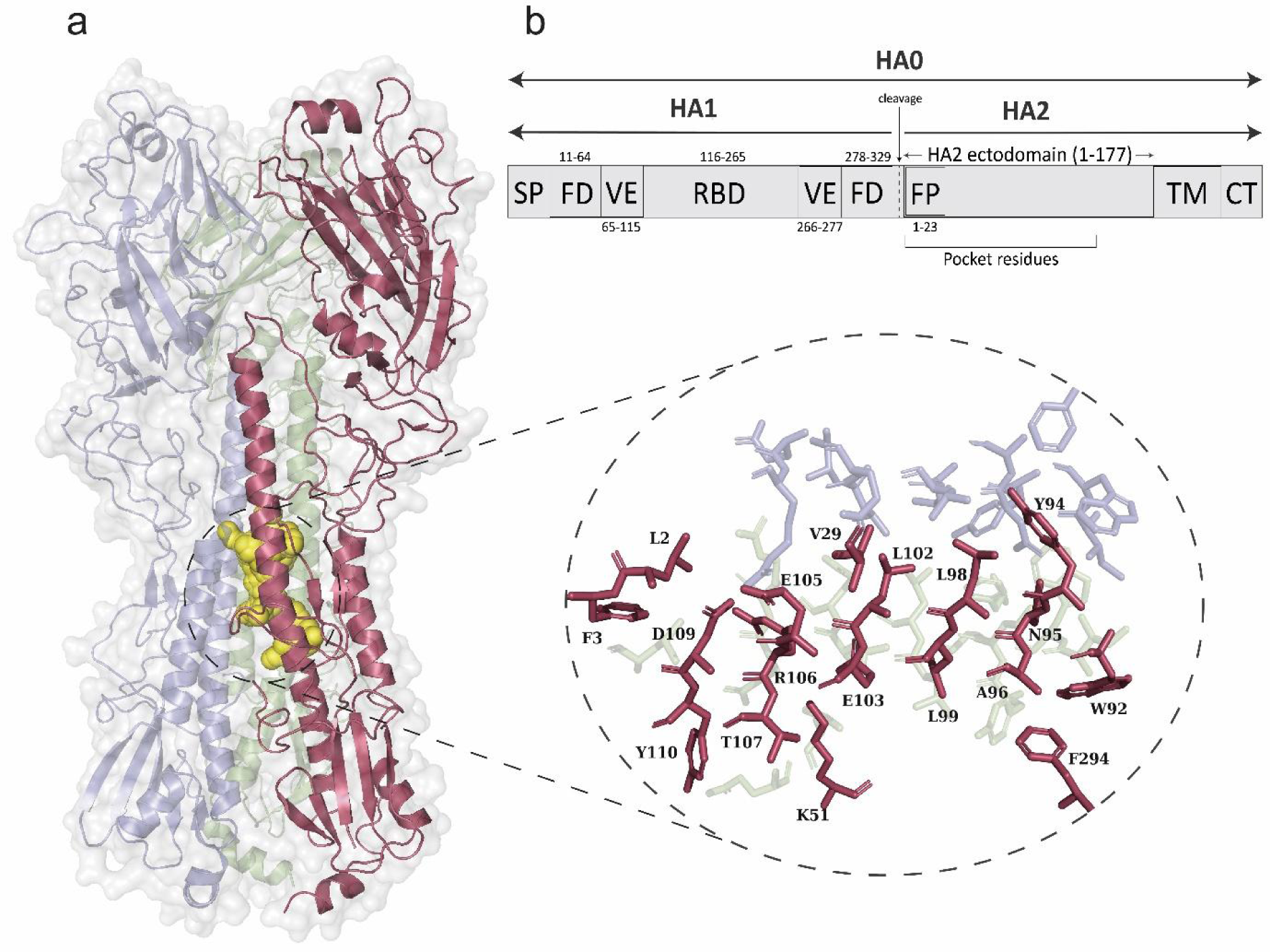
(a) Structural representation of HA trimer of H1N1 (PDB ID: 3LZG). Three protomers are color coded as red, green, and blue respectively. The predicted binding pocket which is sandwiched between three protomers is highlighted as yellow spheres and the residues are shown in an enlarged section. (b) Schematic representation of distinct functional domains of HA protein (SP- Signaling peptide, FD- Fusion domain, VE- Vestigial esterase, FP- Fusion peptide, TM- Transmembrane domain, CT- Cytoplasmic tail)

### Potential Inhibitors of HA identified using Molecular docking and MM-GBSA calculations

A total of 2,471 FDA-approved small molecules were virtually screened against the predicted binding pocket of HA protein and the top-scoring molecules were considered for further analysis (Table 1). It was observed that Econazole exhibited the highest docking score (Glide docking score= −10.454) followed by Butoconazole (−10.448), Miconazole (−9.763), Isoconazole (−9.678), and Tioconazole (−9.312). Noticeably, all the top 5 molecules are imidazole derivatives with broad-spectrum antifungal activity, and Econazole, Butoconazole, and Miconazole are reported to have antibacterial activity (Ivanov et al., 2022; T. Li et al., 2019; Nenoff et al., 2017) (Supplementary table S1). Moreover, Econazole, Butoconazole, Miconazole, and Tioconazole are reported to have anti-influenza activity, although their binding properties remain unclear (An et al., 2014). It is seen that the amino acid residue E-103 of the HA2 is involved in forming a hydrogen bond and salt-bridge interaction in all the 5 top-scoring protein-ligand complexes (Figure 2). Residues Y-94, A-96, L-98, L-99, and L-102 of the HA2 subunit are seen to be involved in hydrophobic interactions with the ligand in these complexes. Polar residue N-95 (HA2) is also observed to interact with the top-scoring ligands. It is vital to take note of the fact that these interacting residues are part of the highly conserved heptad repeat region found close to the fusion peptide (Sriwilaijaroen & Suzuki, 2012; Ghafoori et al., 2023). Such heptad repeats are a common feature observed in amino acid sequences of fusion glycoproteins of several viruses, such as paramyxoviruses, and retroviruses (Chambers et al., 1990).

**Figure 2:**
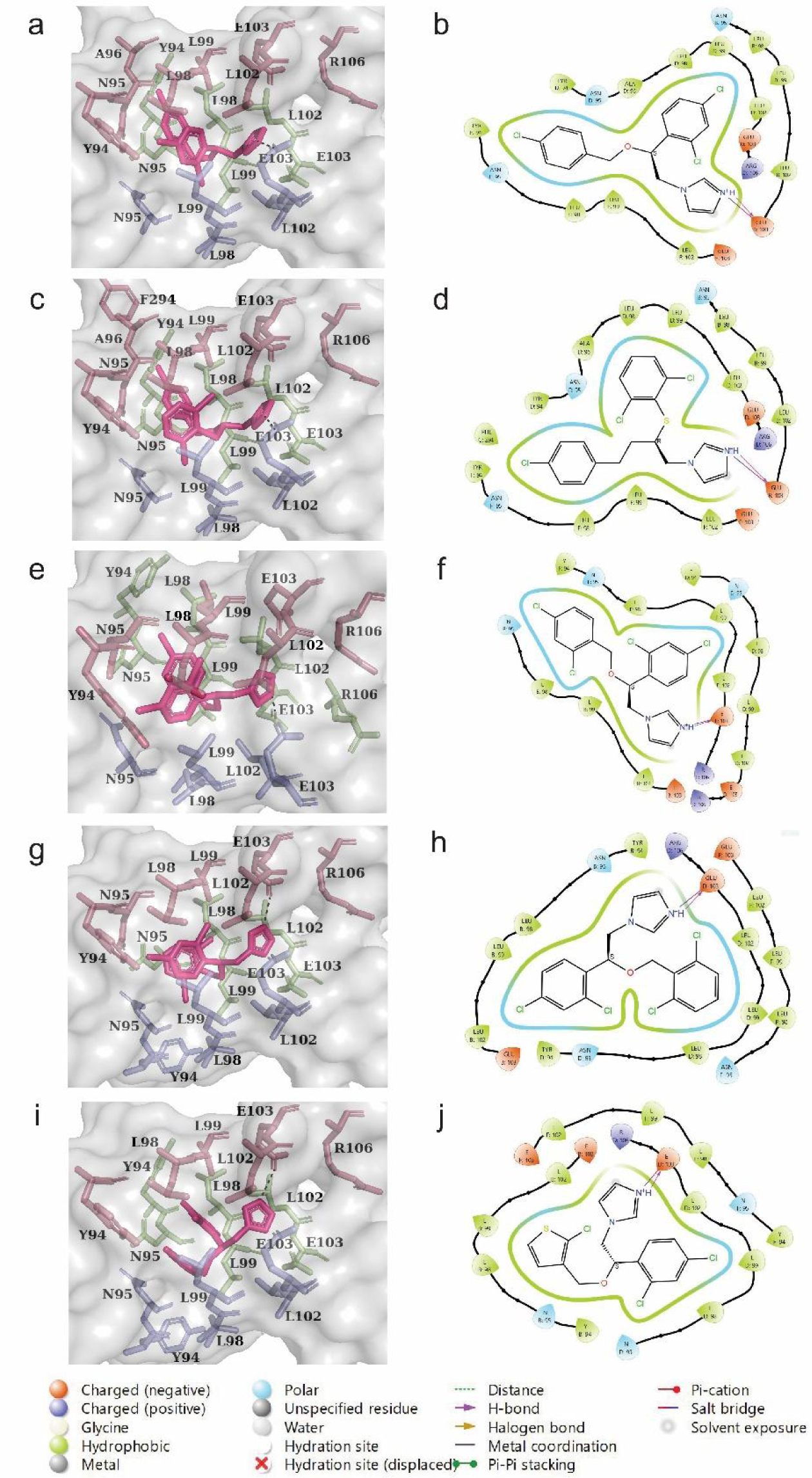
Three-dimensional representation (black dotted line showing H-bonding) and Interaction diagram of top 5 molecules in complex with HA protein showing various bond information. (a, b) Econazole (docking score= −10.45, ΔG bind= −70.38kcal/mol), (c, d) Butoconazole (docking score= −10.45, ΔG bind= −69.56 kcal/mol), (e, f) Miconazole (docking score= −9.76, ΔG bind= −68.44 kcal/mol), (g, h) Isoconazole (docking score= −9.68, ΔG bind= −73.63 kcal/mol), (i, j) Tioconazole (docking score= −9.31, ΔG bind= −64.96 kcal/mol)

**Table 1.** Docking score and MMGBSA ΔG values of 10 top-scoring molecules screened against HA.

| SI No | Drugbank ID | Molecule name | Glide docking score | $\Delta G$ Bind (kcal/mol) | $\Delta G$ Bind Coulomb (kcal/mol) | $\Delta G$ Bind Covalent (kcal/mol) | $\Delta G$ Bind Hbond (kcal/mol) | $\Delta G$ Bind Lipo (kcal/mol) | $\Delta G$ Bind vdW (kcal/mol) |
| --- | --- | --- | --- | --- | --- | --- | --- | --- | --- |
| 1 | DB01127<sup>a</sup> | Econazole | -10.454 | -70.38 | -44.70 | 7.32 | -0.72 | -29.97 | -51.93 |
| 2 | DB00639<sup>a</sup> | Butoconazole | -10.448 | -69.56 | -46.67 | 4.64 | -0.77 | -29.25 | -46.71 |
| 3 | DB01110<sup>a</sup> | Miconazole | -9.763 | -68.44 | -43.35 | 6.79 | -0.03 | -31.42 | -50.24 |
| 4 | DB08943 | Isoconazole | -9.678 | -73.63 | -55.28 | 3.52 | -0.60 | -28.24 | -44.43 |
| 5 | DB01007<sup>a</sup> | Tioconazole | -9.312 | -64.96 | -37.19 | 4.46 | -0.78 | -25.81 | -46.13 |
| 6 | DB00557<sup>a</sup> | Hydroxyzine | -8.513 | -66.07 | -50.37 | 5.29 | -0.86 | -32.8 | -43.69 |
| 7 | DB01114<sup>a</sup> | Chlorpheniramine | -8.391 | -63.44 | -30.77 | 1.64 | -0.42 | -26.71 | -44.57 |
| 8 | DB01624 | Zuclopenthixol | -8.193 | -64.33 | -33.66 | 7.08 | -0.57 | -36.64 | -48.20 |
| 9 | DB01551 | Dihydrocodeine | -7.860 | -63.38 | -34.3 | 4.95 | -0.72 | -24.01 | -43.13 |
| 10 | DB08936 | Chlorcyclizine | -7.723 | -62.07 | -35.66 | 1.22 | -0.04 | -28.23 | -43.55 |

<sup>a</sup> represents molecules that have been reported to have anti-influenza activity (An et al. 2014; Xu et al. 2018)

A deeper insight into the ligand binding energies can be obtained by the MMGBSA calculations. HA complexed with Isoconazole showed the highest binding free energy (−73.63 kcal/mol), followed by that of Econazole (−70.38 kcal/mol), Butoconazole (−69.56 kcal/mol), Miconazole (−68.44 kcal/mol) and Tioconazole (−64.96 kcal/mol). It is observed that the most contributions towards the binding energy are by the coulomb energy, Van der Waals energy, lipophilic energy, and H-bond energy (Table 1). Thus, the molecular docking analysis and MMGBSA results indicate that coulombic/electrostatic interactions, lipophilic/hydrophobic interactions, and hydrogen bonding could be the driving forces for the protein-ligand binding in the 5 top-scoring HA-ligand complexes.

### Molecular dynamics simulation of HA complexed with potential inhibitors

The time-dependent binding stability of the top scoring 5 molecules was validated through MD simulations over 100ns. The results were analyzed in terms of root mean square deviation (RMSD), root mean square fluctuation (RMSF), and protein-ligand contacts. For the compounds Econazole, Butoconazole, and Isoconazole the protein RMSD stabilizes after the initial 20 ns, and later variations of the order of ∼2 Å are observed and the average protein RMSD values during the simulations are 3.39 Å, 3.27 Å, and 3.18 Å, respectively (Figure 3 (a), (b), (d)). For Miconazole and Tioconazole, the protein RMSD stabilizes after 30 ns and the variations after that are of ∼2 Å, while the average RMSD throughout simulation is 3.18 Å and 3.27 Å for the two complexes (Figure 3 (c), (e)). The ligand RMSD for Econazole, Butoconazole, Miconazole, and Isoconazole stabilizes after initial 10-20 ns. Post stabilization, the variation in ligand RMSD observed was around 2-3 Å. The RMSF plots of the top 5 complexes show that average RMSF values are 1.37 Å, 1.42 Å, 1.48 Å, 1.22 Å, and 1.65 Å. Higher fluctuations correspond to the N and C-terminal residues of the two subunits of each protomer. Almost all the interacting residues showed RMSF values less than 1 Å (indicated by green lines, Supplemetray figure S1).

**Figure 3:**
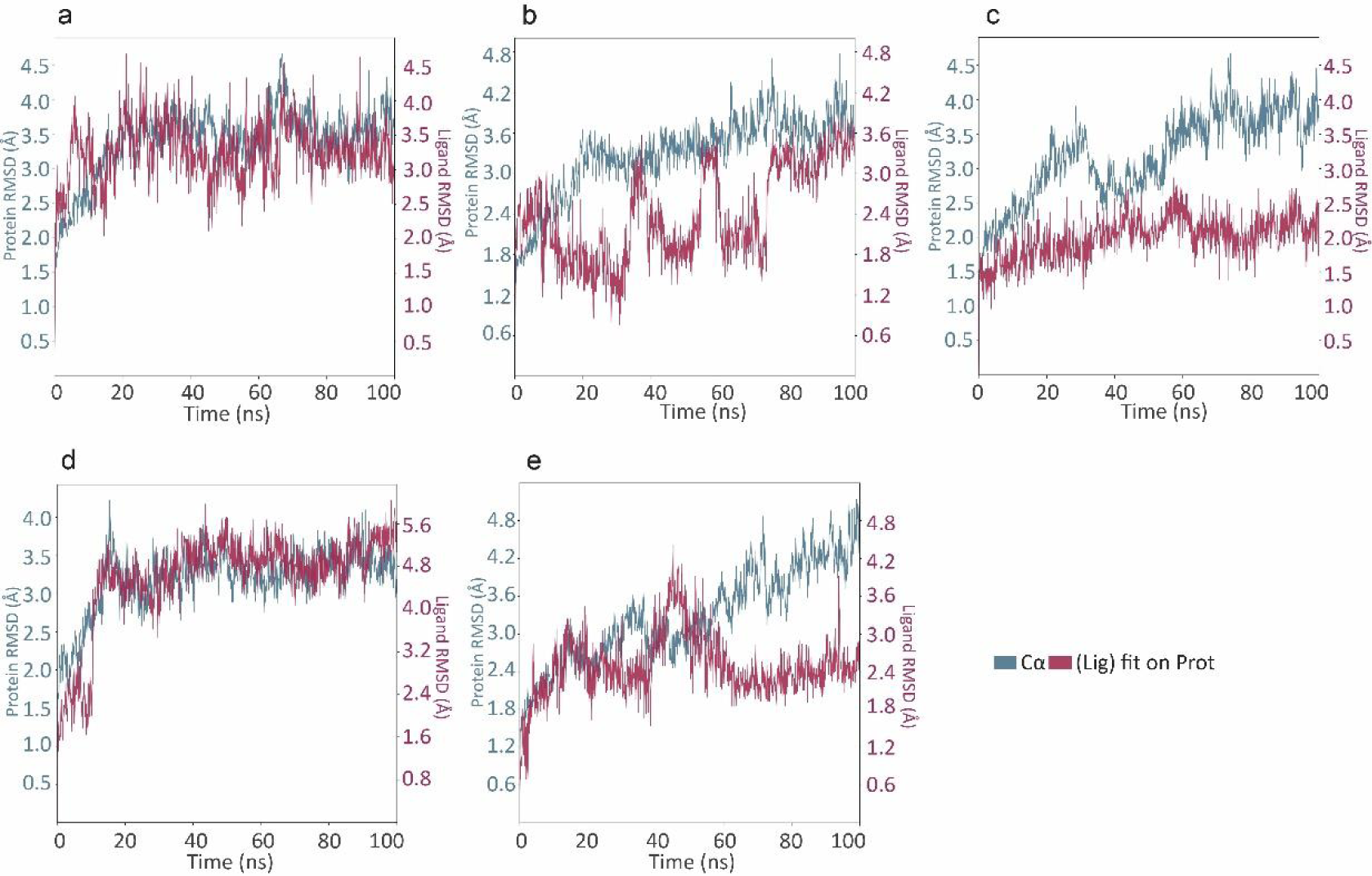
Plots representing RMSD of top scoring (five) ligand-protein complexes simulated for 100ns. (a) HA-Econazole complex, (b) HA-Butoconazole complex, (c) HA-Miconazole complex, (d) HA-Isoconazole complex, and (e) HA-Tioconazole complex

Our MD simulation analysis shows that Butoconazole has the highest number of interacting amino acid residues (18), followed by Econazole (16), Tioconazole (16), Isoconazole (15), and Miconazole (13) (Figure 4). E-103 of HA2 subunit in each protomer is an important residue that maintains hydrogen bond, water bridges, and ionic interactions in all 5 top-scoring protein-ligand complexes. In Miconazole, N-95 is also involved in forming hydrogen bonding. Residues Y-94, L-98, L-99, and L-102 of the HA2 subunit are important in maintaining hydrophobic bonds in all the 5 top-scoring complexes. V-29 of the HA1 subunit is also involved in hydrophobic interaction in the HA-Tioconazole complex.

**Figure 4:**
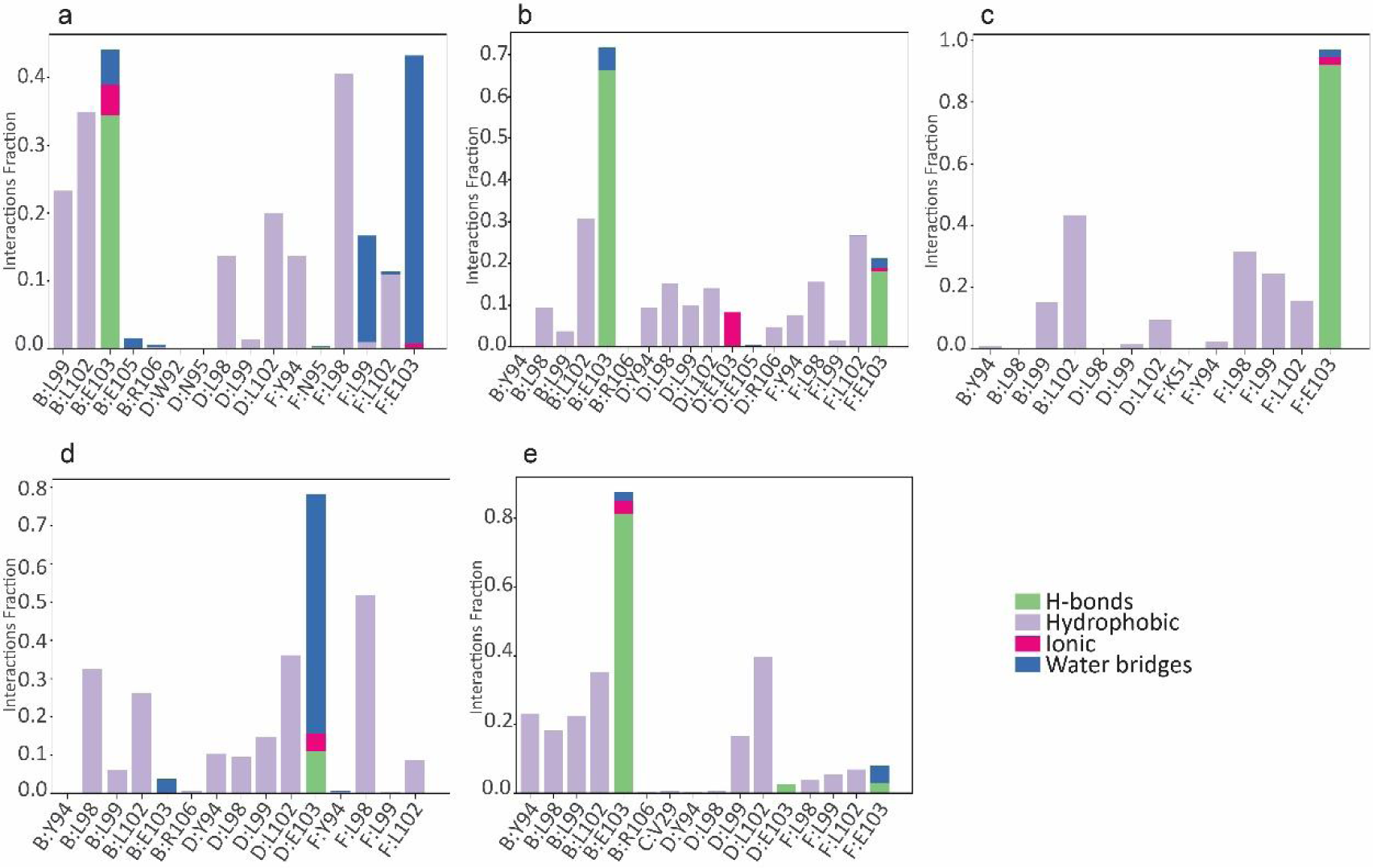
Interaction fraction and interacting protein residues of top scoring (five) ligand-protein complexes simulated for 100 ns. (a) HA-Econazole complex, (b) HA-Butoconazole complex, (c) HA-Miconazole complex, (d) HA-Isoconazole complex, and (e) HA-Tioconazole complex

### Trajectory analysis of HA complexed with potential inhibitors

After the 100 ns simulations, Principal component analysis (PCA) was done for the trajectories of the top 5 HA-ligand complexes and the apo-HA protein (unbound form). The principal components PC1 and PC2 captured 41.317% aggregate motion in HA-econazole complex, 44.050% aggregate motion in HA-butoconazole complex, 41.608% aggregate motion in HA-miconazole complex, 38.999% aggregate motion in HA-isoconazole complex, and 52.279% aggregate motion in HA-tioconazole complex. The PCA plot shows that the collective motions of the protein-ligand complexes have not deviated from that of apo-HA and occupy less conformational space, indicating stabilization of the trimeric pre-fusion structure of the HA protein (Figure 5).

**Figure 5:**
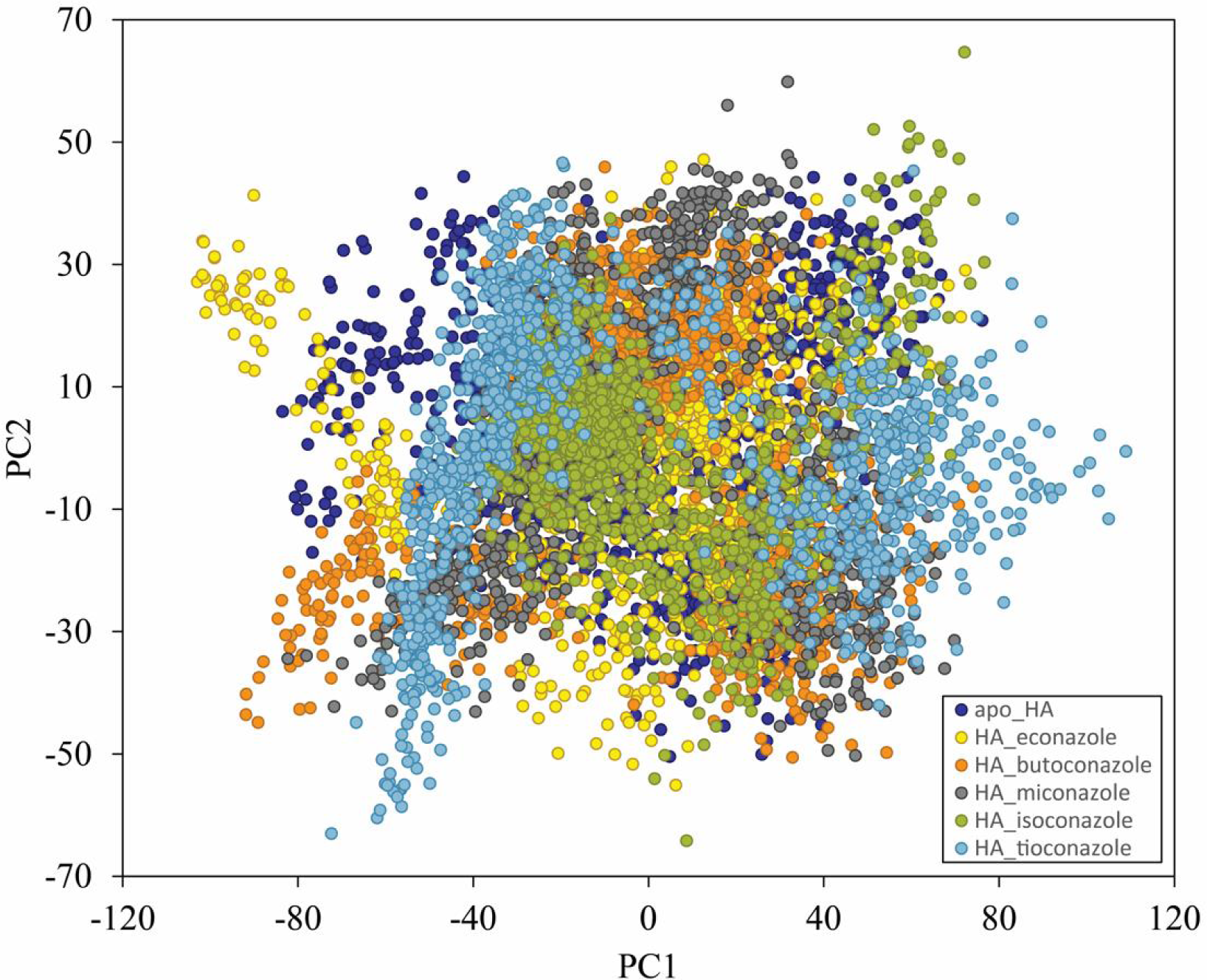
Principal component analysis of MD simulation trajectories for apo-HA protein (dark blue) and top-scoring (five) HA complexes with Econazole (yellow), Butoconazole (orange), Miconazole (grey), Isoconazole (green), and Tioconazole (blue)

2-D contour maps of Free Energy Landscape (FEL) corresponding to the top 5 HA-ligand complexes were generated using RMSD and RG (radius of gyration) as the reaction coordinates in order to determine the global minimum energy conformation during the 100 ns simulations. Red colour represents a higher energy state, while blue indicates a lower energy state. The results indicate that the lowest free energy state (deep blue colour) was achieved by the protein-ligand complexes (Figure 6). The size and shape of this area revealed the stability of complexes. A single centralized and compact global minimum energy in the case of the HA-Econazole complex indicates minimum structural changes. The presence of 2 or more minimum energy states within the low free energy area in the case of the other four protein-ligand complexes indicates multiple stable states which could be due to the presence of multiple binding modes as seen during the trajectory visualization. Thus, the PCA and FEL results help explain the ligand-induced conformational changes indicated by the RMSD and RMSF plots.

**Figure 6:**
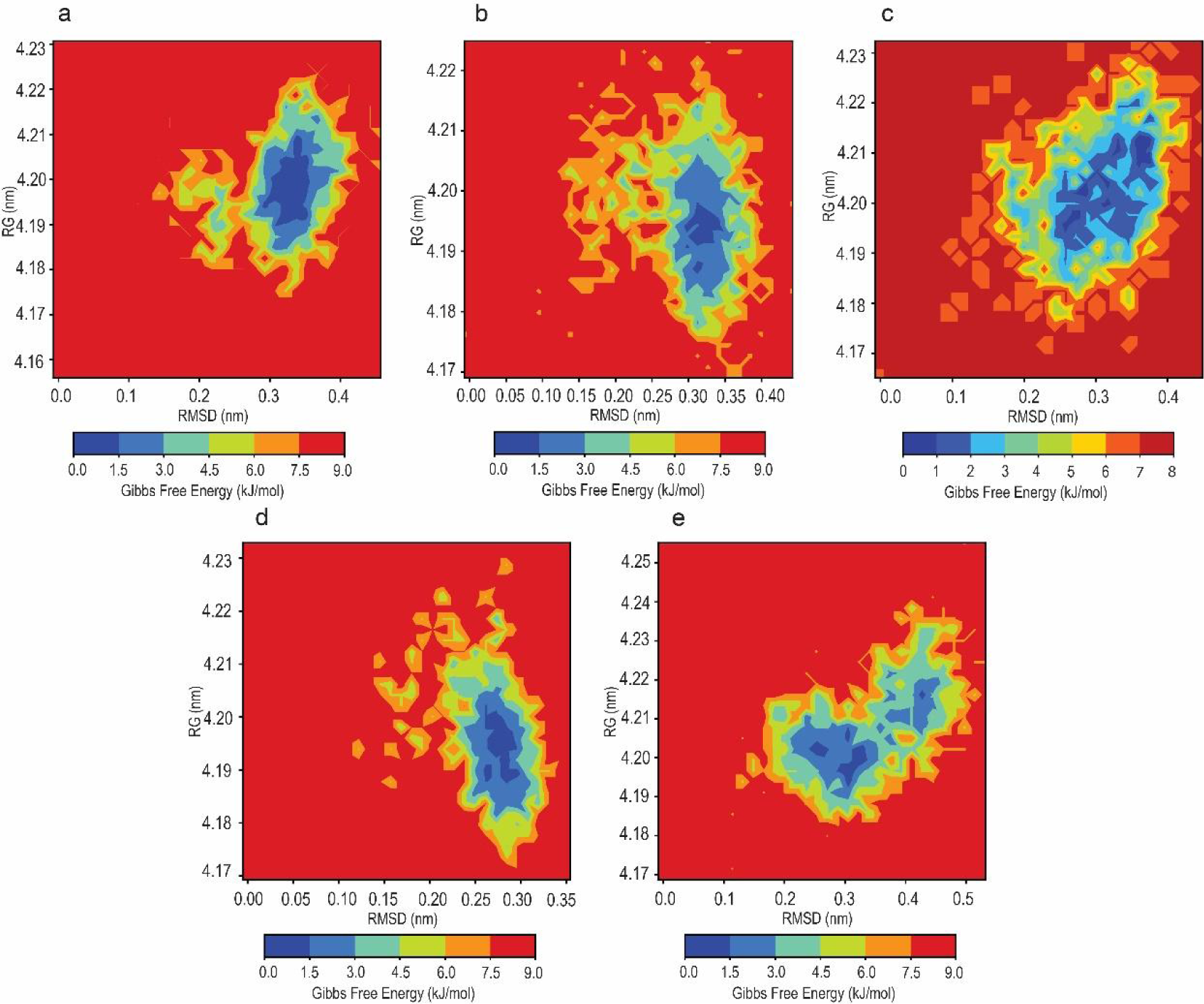
Two-dimensional contour map of Free Energy Landscape (FEL) of protein-ligand complexes as a function of RMSD and RG. (a) FEL of HA-Econazole, (b) FEL of HA-Butoconazole, (c) FEL of HA-Miconazole, (d) FEL of HA-Isoconazole, and (e) FEL of HA-Tioconazole

### Ligand Binding Site of NA

The substrate binding site of NA near the 150-loop, where the co-crystal ligand oseltamivir carboxylate was bound, was considered for molecular docking. A SiteMap prediction was also carried out without the co-crystal ligand and the top-scoring site (site score =1.07) coincides with the chosen site. The residues forming the binding pocket and within 4 Å vicinity of the co-crystal ligand are R-118, E-119, D-151, R-152, W-178, I-222, R-224, S-246, E-276, E-277, R-292, N-294, R-371, and Y-406 (Figure 7).

**Figure 7:**
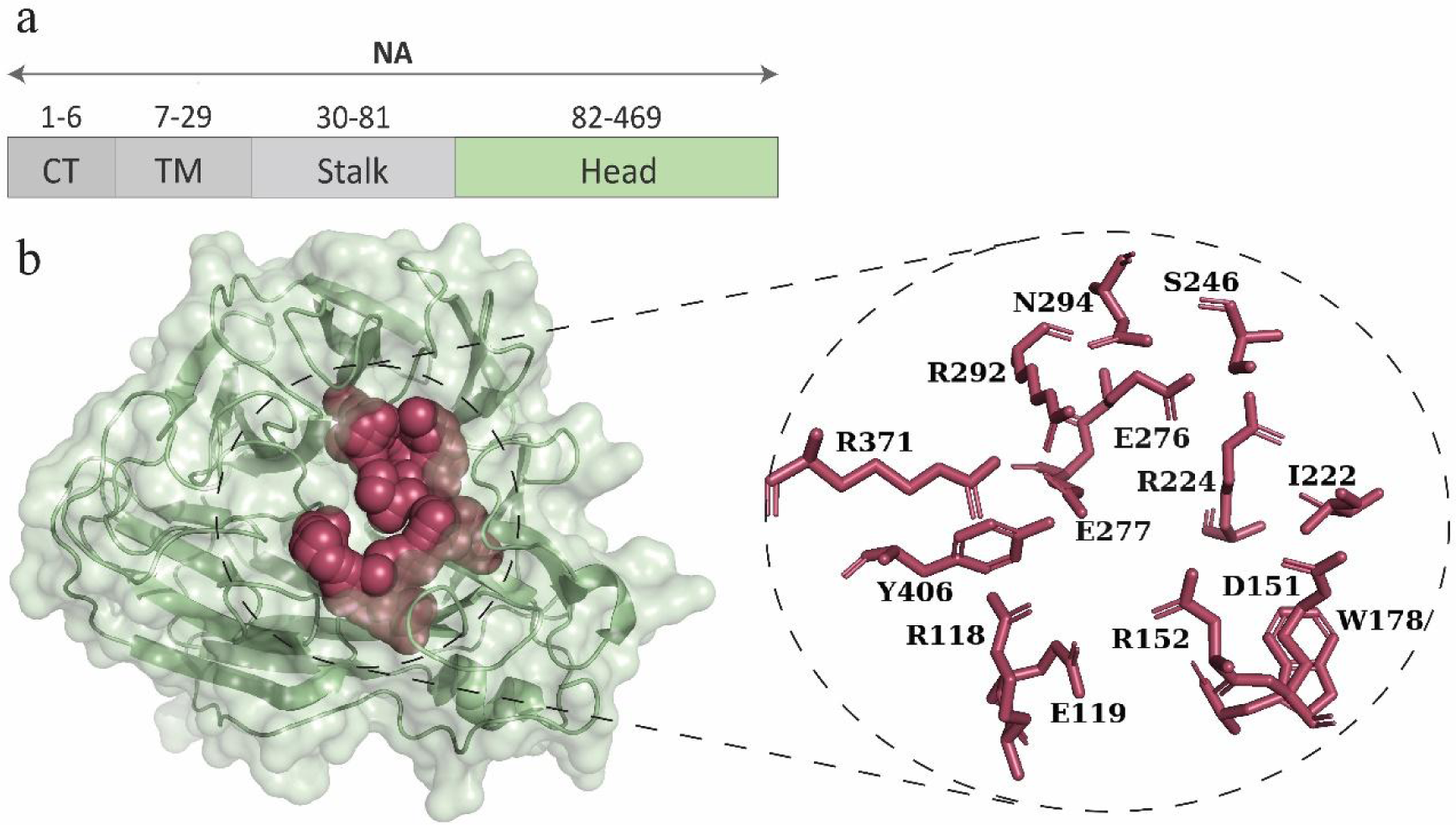
(a) Schematic representation of distinct functional domains of H1N1 NA protein. (b) Structural representation of the head domain of NA monomeric subunit (PDB ID: 3TI6). The binding site region is highlighted as red spheres and amino acid residues are shown in the enlarged section

### Potential Inhibitors of NA identified using Molecular docking and MM-GBSA calculations

After virtually screening 2,471 FDA-approved small molecules against the substrate binding site of NA, the top-scoring NA-ligand complexes were considered for further analysis (Table 2). It was observed that the molecule Acarbose has the highest Glide docking score of −10.93, followed by Rutin (−10.18), Paromomycin (−9.61), Idarubicin (−7.31), and Dabigatran (−7.05). The top-scoring molecule Acarbose, an oral drug prescribed to diabetic patients, has been reported to have an inhibitory effect on Enterovirus 71 and has also shown promising results in in-silico drug screening studies against SARS-CoV-2 (Feng et al., 2020; Kumar et al., 2020; Sundar et al., 2021). Among these, molecules Rutin (a flavonoid) and Paromomycin (a broad-spectrum aminoglycoside antibiotic) have been reported to have anti-influenza activity through immunomodulatory effect and inhibiting the virus promoter (vRNA) respectively (Kim et al., 2012; Ling et al., 2020).

**Table 2.**
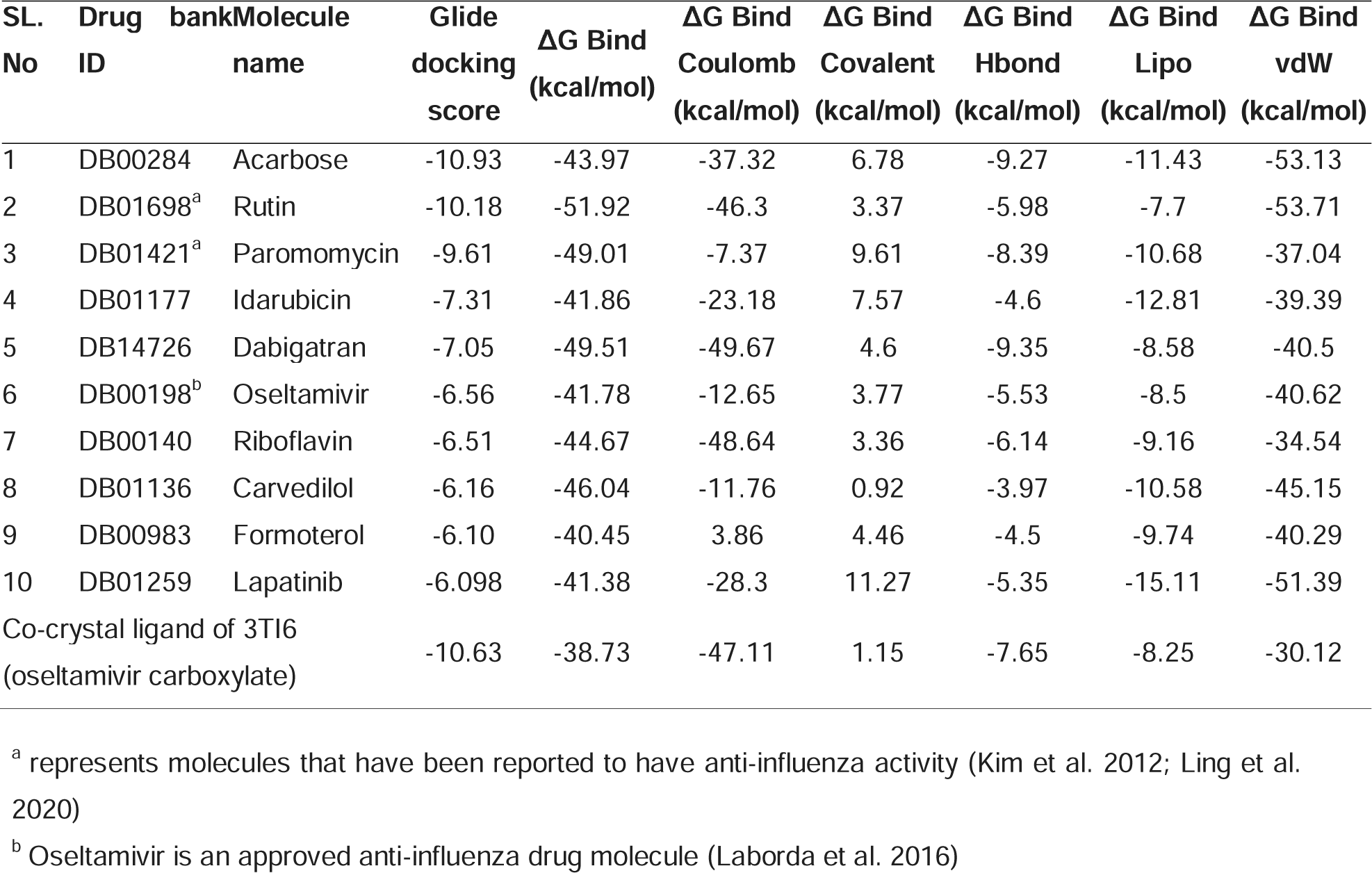
Docking score and MMGBSA ΔG values of 10 top-scoring molecules screened against NA.

A comparison of the binding free energy values in the 5 top-scoring NA-ligand complexes, obtained from the MMGBSA calculations, shows that the NA-Rutin complex has the highest binding energy of −51.92 kcal/mol, followed by Dabigatran −49.51 kcal/mol, Paromomycin −49.01 kcal/mol, Acarbose −43.97 kcal/mol, and Idarubicin −41.86 kcal/mol. Most of the contributions towards the binding free energy in the top 5 complexes are by the coulomb energy, Van der Waals energy, lipophilic energy, and H-bond energy. Residue R-118 was seen to be interacting with Acarbose and Dabigatran via H-bond. E-119 interacted with Paromomycin through H-bonds, and with Idarubicin and Dabigatran through water-bridges. D-151 formed 3 H-bonds in Acarbose, one H-bond with Rutin, 2 H-bonds and 2 water bridge interactions with Paromomycin, and a H-bond and a water bridge interaction with Dabigatran. R-152 was seen to be interacting with Acarbose and Rutin via H-bonding. W-178 interacted with Acarbose, Paromomycin, and Idarubicin through H-bonding as well. Residue E-227 was seen to be forming H-bond in all the 5 protein-ligand complexes. It also formed a water bridge interaction with Paromomycin and Idarubicin. Residue E-277 interacted with Rutin and Paromomycin through H-bonding. It also interacted with Acarbose and Idarubicin via water bridge interaction. Residue R-371 is also important in maintaining H-bond with Acarbose, Rutin, Idarubicin, and Dabigatran (Figure 8). It is noteworthy that the residues R-118, D-151, R-152, and R-371 are highly conserved residues that form the inner shell of the active site and are known to directly interact with the sialic acid substrates. Residues E-119, W-178, E-227, and E-277, though do not directly interact with the substrate, they form the outer shell of the active site and are referred to as the framework residues (Shtyrya et al., 2009).

**Figure 8:**
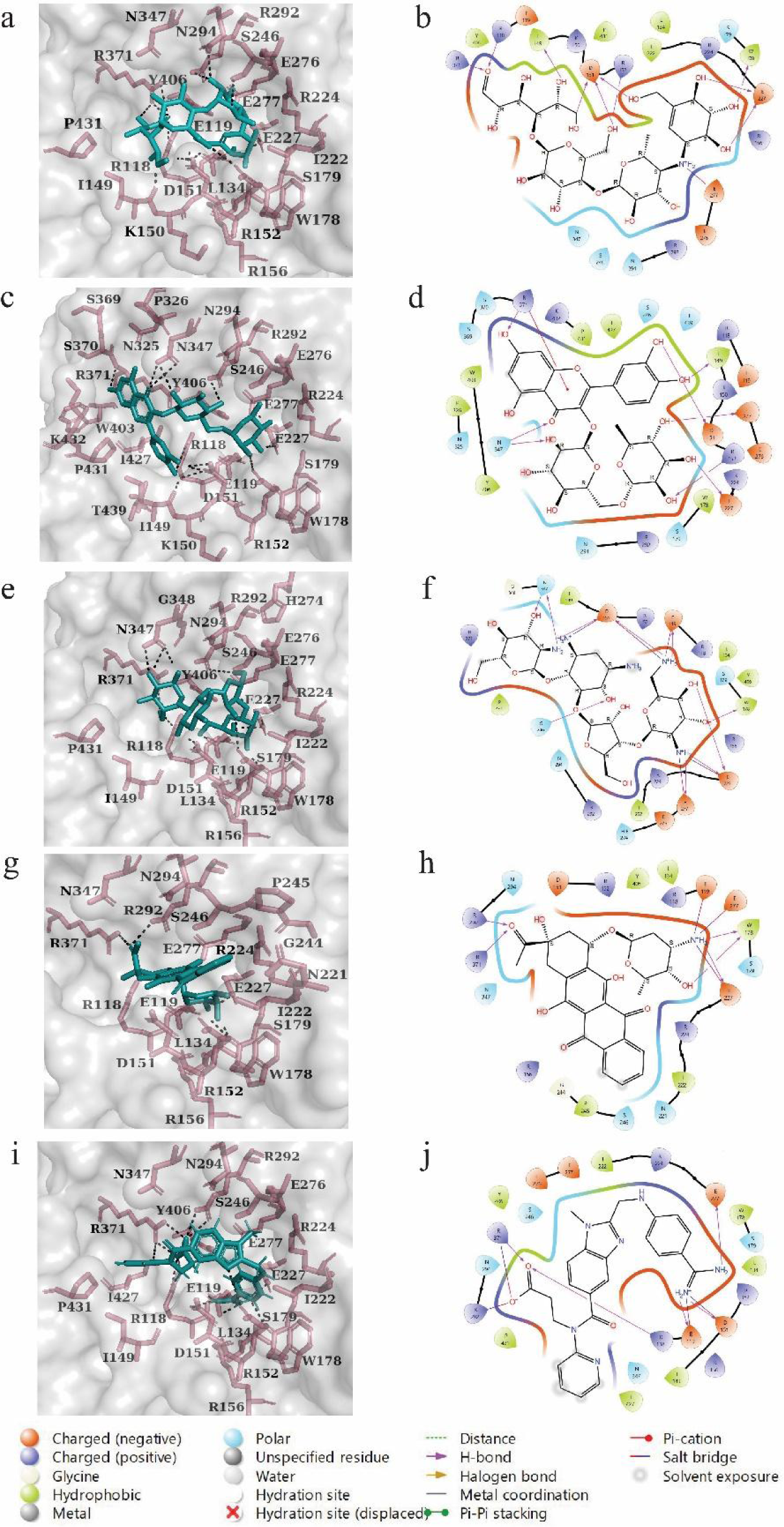
Three-dimensional representation (black dotted lines showing H-bonding) and Interaction diagram of top 5 molecules in complex with NA protein showing various bond information. (a, b) Acarbose (docking score= −10.93, ΔG bind= −43.97 kcal/mol), (c, d) Rutin (docking score= −10.18, ΔG bind= −51.92 kcal/mol), (e, f) Paromomycin (docking score= −9.61, ΔG bind= −49.01 kcal/mol), (g, h) Idarubicin (docking score= −7.31, ΔG bind= −41.86 kcal/mol), (i, j) Dabigatran (docking score= −7.05, ΔG bind= −49.51 kcal/mol)

### Molecular dynamics simulation of NA complexed with potential inhibitors

The RMSD plots for NA-ligand complexes of the 5 top scoring molecules showed that the protein RMSD stabilizes after the initial 10-20 ns. The average protein RMSD values were 2.25 Å for Acarbose, 2.8 Å for Rutin, 1.66 Å for Paromomycin, 2.24 Å Idarubicin, and 2.13 Å for Dabigatran. The ligand RMSD for Paromomycin and Idarubicin stabilizes after 10-20 ns, whereas it takes around 40 ns for the stabilization of Acarbose and Rutin, and 65 ns for Dabigatran (Figure 9). The average RMSF values for the 5 complexes were in the range of 0.80-0.95 Å. Higher fluctuations were shown by the C-terminal residues. The interacting residues showed RMSF of ∼2 Å and lower, except in the case of Dabigatran where the interacting residue I-149 showed a RMSF of 3 Å (Supplementary figure S2). From the MD simulation analysis it was seen that Rutin had the highest number of interacting residues (45), followed by Acarbose and Paromomycin (27), Dabigatran (16), and Idarubicin (15). Charged amino acids like E-119, D-151, R-152, E-227, and E-277 are involved in binding via hydrogen bonds, water bridge interactions, and ionic interactions. Interactions in the case of the NA-dabigatran complex were seen to be lost gradually (Figure 10). Thus, our MD simulation analysis showed stable NA-ligand complex formation in the case of Acarbose, Rutin, Paromomycin, and Idarubicin.

**Figure 9:**
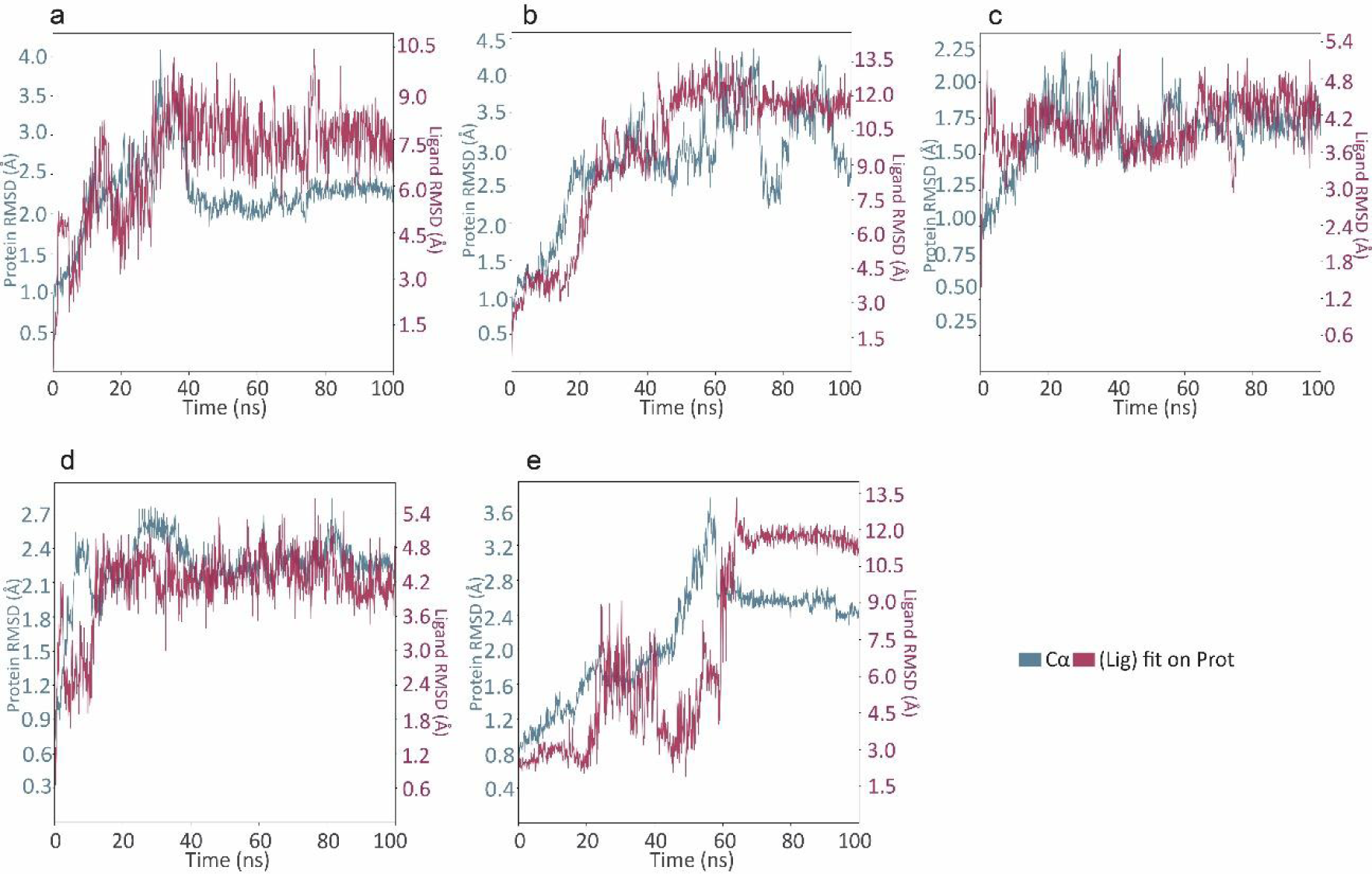
Plots representing RMSD of top scoring (five) ligand-protein complexes simulated for 100ns. (a) NA-Acarbose complex, (b) NA-Rutin complex, (c) NA-Paromomycin complex, (d) NA-Idarubicin complex, and (e) NA-Dabigatran complex

**Figure 10:**
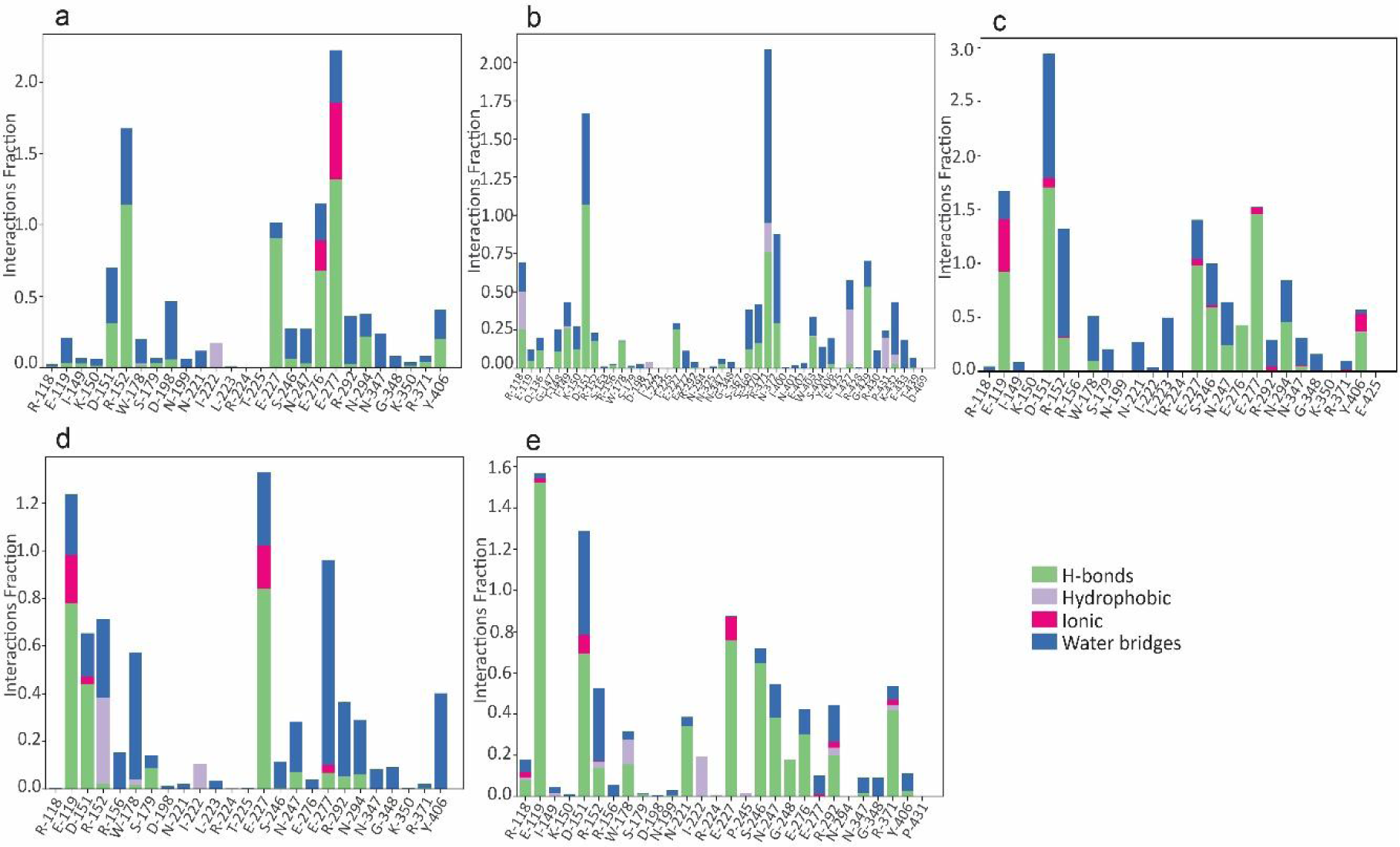
Interaction fraction and interacting protein residues of top scoring (five) ligand-protein complexes simulated for 100 ns. (a) NA-Acarbose complex, (b) NA-Rutin complex, (c) NA-Paromomycin complex, (d) NA-Idarubicin complex, and (e) NA-Dabigatran complex

### Trajectory analysis of NA complexed with potential inhibitors

Principal component analysis (PCA) of the trajectories, obtained after the 100 ns MD simulation, was done for unbound NA protein (apo form), and the top 5 NA-ligand complexes. The principal components PC1 and PC2 captured 51.991% aggregate motion in NA-acarbose complex, 63.640% aggregate motion in NA-rutin complex, 40.246% aggregate motion in NA-paromomycin complex, 47.412% aggregate motion in NA-idarubicin complex, and 59.826% aggregate motion in NA-dabigatran complex. The PCA scatter plot showed that the collective motions of the protein-ligand complexes have not deviated from that of apo-NA except for that of Rutin and Dabigatran and occupy less conformational space, indicating stable binding (Figure 11). Although protein RMSD and PCA analysis of Rutin showed higher deviation, the trajectory visualization revealed it to be binding in the substrate binding site of NA throughout the simulation which may be attributed to its highest number of protein-ligand contacts.

**Figure 11:**
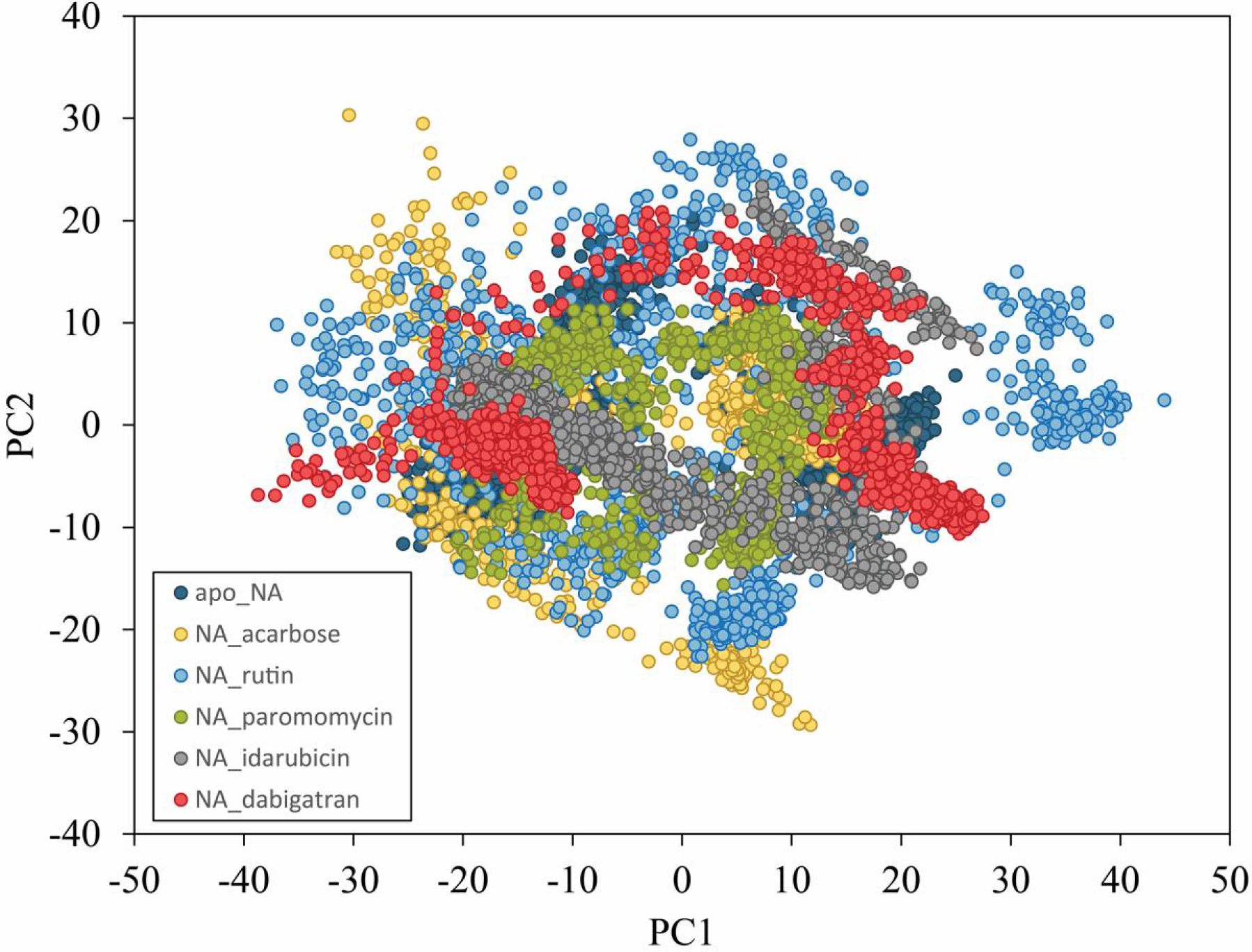
Principal component analysis of MD simulation trajectories for apo-NA protein (dark blue) and five top-scoring NA complexes with acarbose (yellow), rutin (blue), paromomycin (green), idarubicin (grey), and dabigatran (red)

In order to determine the energy minima of the landscape of the top 5 NA-ligand complexes, we constructed the FEL as a function of RMSD and RG values during the 100 ns simulations. All the protein-ligand complexes were seen to be converging towards the deepest blue region, indicating the attainment of the lowest free energy state (Figure 12). This clearly illustrates the stability of these complexes.

**Figure 12:**
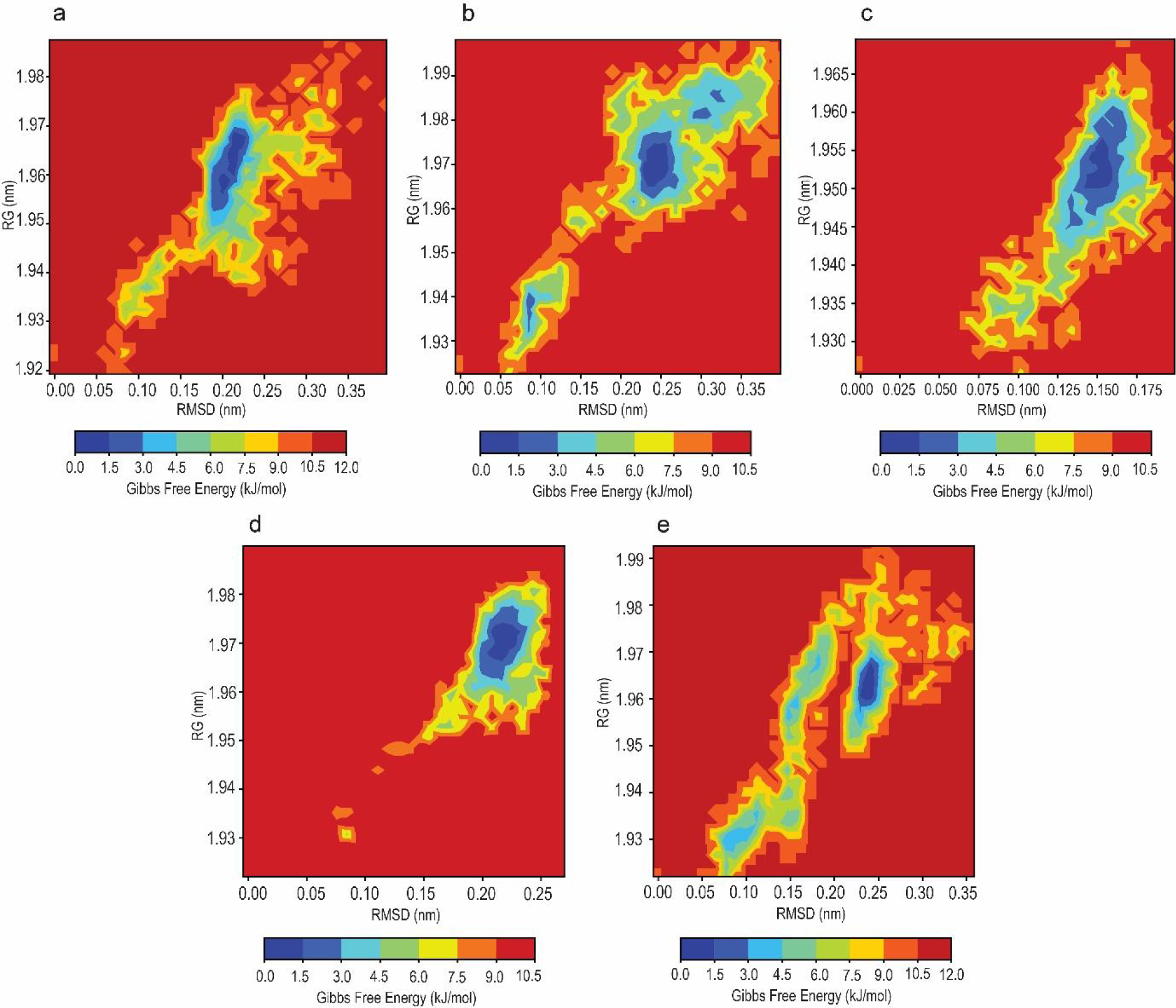
Principal component analysis of MD simulation trajectories for apo-NA protein (dark blue) and five top-scoring NA complexes with acarbose (yellow), rutin (blue), paromomycin (green), idarubicin (grey), and dabigatran (red)

## Conclusions

Our molecular docking analysis, followed by MM-GBSA calculations revealed FDA-approved drug molecules to be potentially binding to HA and NA surface proteins of H1N1. Further analysis of the 100 ns MD simulations comparing the protein-ligand complexes with the apo-forms of the proteins showed the stability of 9 molecules. Our results show that the binding of Econazole, Butoconazole, Miconazole, Isoconazole, and Tioconazole form stable complexes with the pre-fusion trimeric structure of HA. In the case of NA-ligand complexes, we observed stable binding of Acarbose, Rutin, Paromomycin, and Idarubicin. Interestingly, six out of the nine molecules that exhibited strong binding to HA and NA in our study have been previously reported to have anti-influenza activity. This finding not only reinforces the reliability of our virtual screening approach but also highlights the potential of repurposing existing drugs with known antiviral properties. Our study provides valuable insights into the binding sites and interactions of small molecules with the surface proteins of H1N1 and also emphasizes the importance of targeting these proteins as an effective strategy to combat influenza A viruses. By interfering with crucial steps in the viral life cycle, drugs targeting HA and NA have the potential to halt the viral spread and reduce the severity of infections. Further, we hypothesize that the binding of the small molecules at the vicinity of the fusion peptide and fusion domain will inhibit the conformational change that is essential for successful fusion, and in the case of NA, binding of the small molecules blocks the substrate binding site near the 150-loop region of NA and can potentially inhibit the protein. This can be further validated by in-vitro and in-vivo experiments.

## Acknowledgements

The authors wish to thank Prof. Chandrabhas Narayana, the Director, RGCB for providing intramural support and computational infrastructure. We also thank Dr. Sivakumar K.C and Dr. Jamshaid Ali, Bioinformatics Facility, RGCB for useful discussions.

## Author Contributions

SB, RP, SNS: Methodology, Validation, Formal analysis, Investigation, Data curation, Writing—original draft, Visualization; SB, RP, LVS, RC, KN, SNS: Validation, Writing—review & editing; SNS: Conceptualization, Validation, Project administration, Resources, Funding acquisition, review & editing, supervision.

