## Supplementary material for "Structure-based virtual screening and Molecular Dynamic Simulations identified FDA-approved molecules as potential inhibitors against the surface proteins of H1N1": Supplemtary Files

**Supplementary file**

**Supplementary figure S1** Plots representing RMSF (with residue index) of top scoring (five) ligand-protein complexes simulated for 100ns. (a) HA-Econazole complex, (b) HA-Butoconazole complex, (c) HA-Miconazole complex, (d) HA-Isoconazole complex, and (e) HA-Tioconazole complex

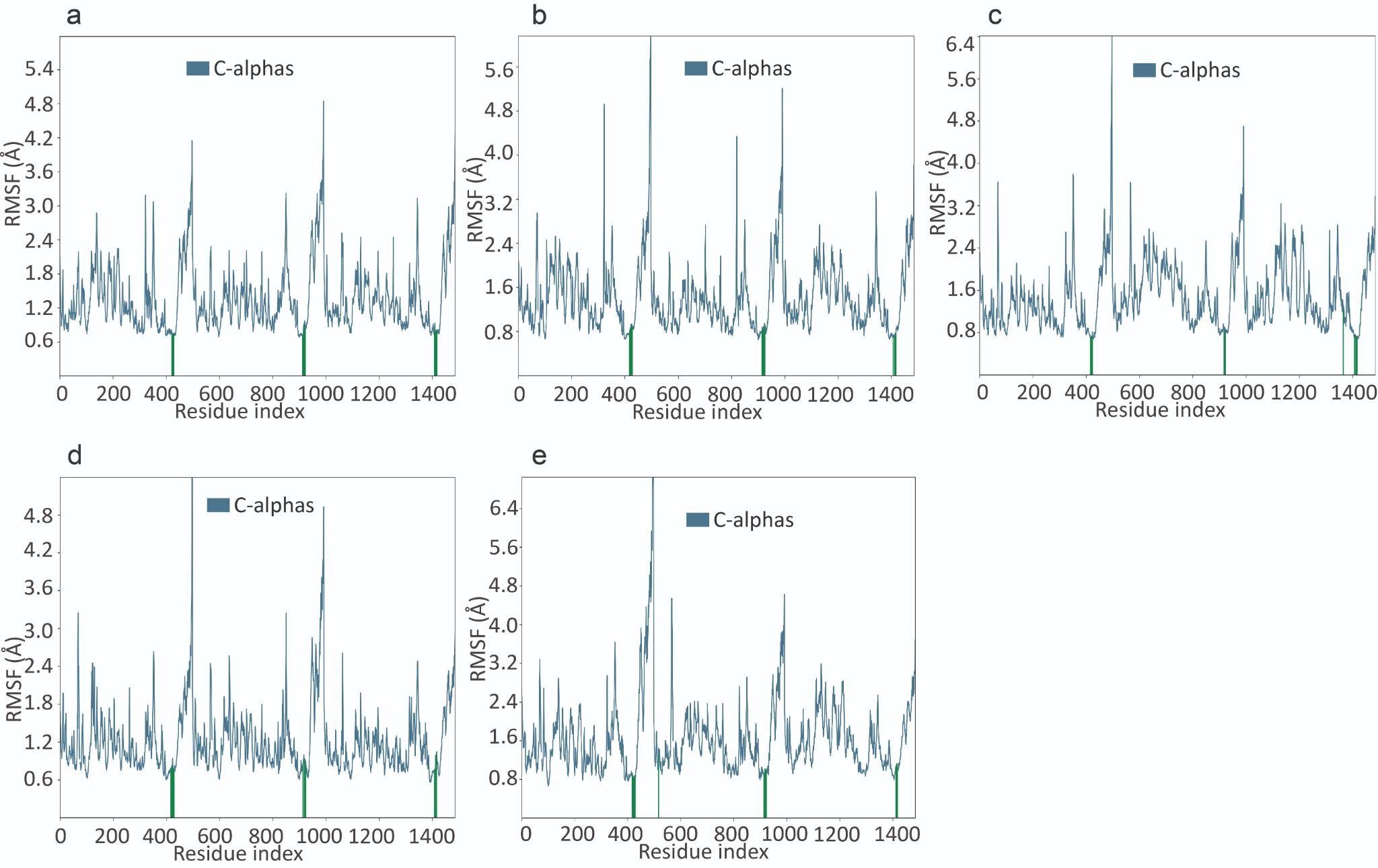

**Supplementary figure S2** Plots representing RMSF (with residue index) of top scoring (five) ligand-protein complexes simulated for 100ns. (a) NA-Acarbose complex, (b) NA-Rutin complex, (c) NA-Paromomycin complex, (d) NA-Idarubicin complex, and (e) NA-Dabigatran complex

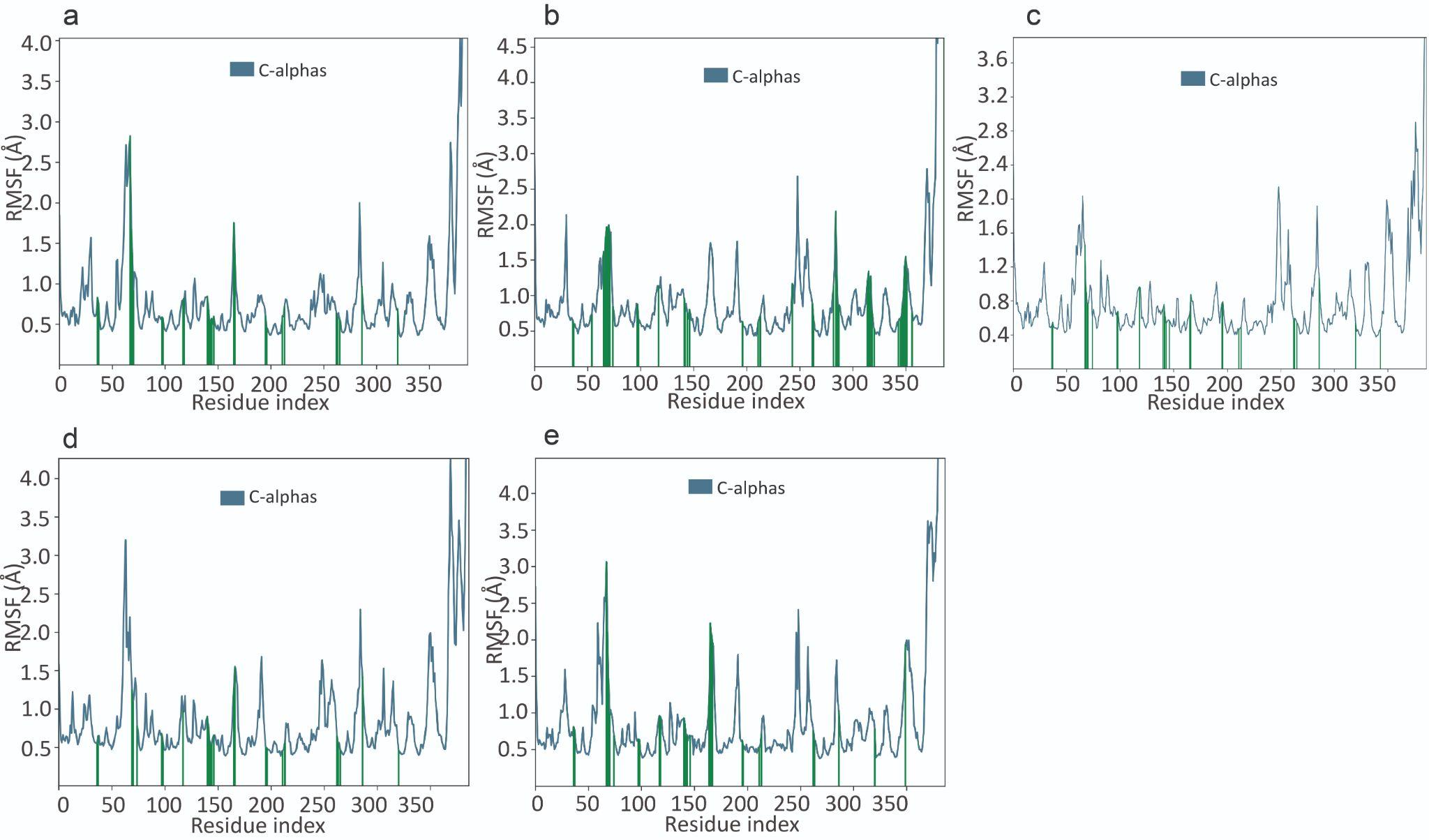

**Supplementary table S1** Two-dimensional structures of 10 top-scoring molecules docked with HA

| **SL. No** | **Drugbank ID** | **Molecule name** | **2-D Structure** | **Previously reported functions** |
| --- | --- | --- | --- | --- |
| 1 | DB01127 | Econazole | 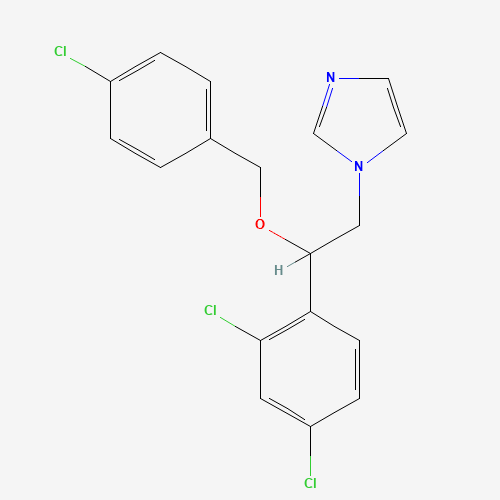 | Antifungal drug with antibacterial and anti-influenza activity (An et al. 2014; Qiu et al. 2017) |
| 2 | DB00639 | Butoconazole | 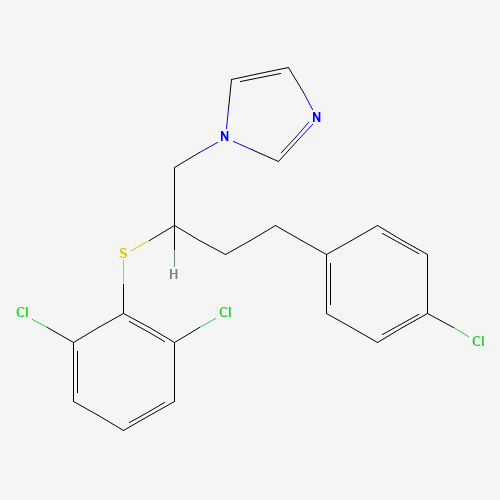 | Antifungal drug with anti-influenza activity (An et al. 2014) |
| 3 | DB01110 | Miconazole | 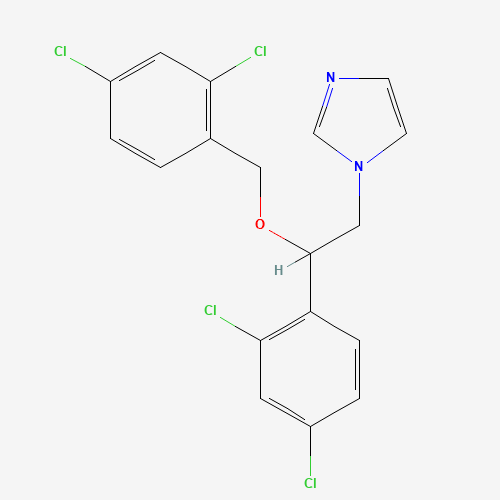 | Antifungal drug with antibacterial and anti-influenza activity (An et al. 2014; Nenoff et al. 2017) |
| 4 | DB08943 | Isoconazole | 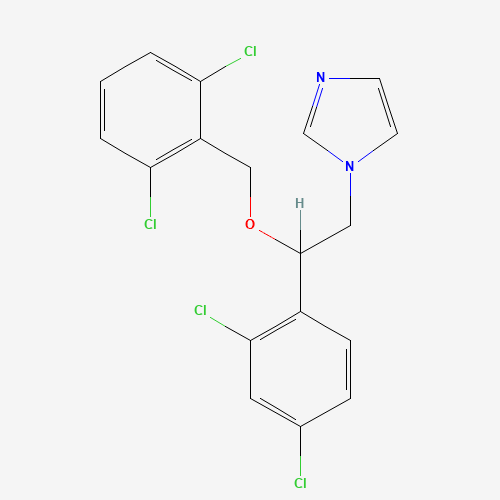 | Antifungal drug with antibacterial activity (Czaika et al. 2013; An et al. 2014) |
| 5 | DB01007 | Tioconazole | 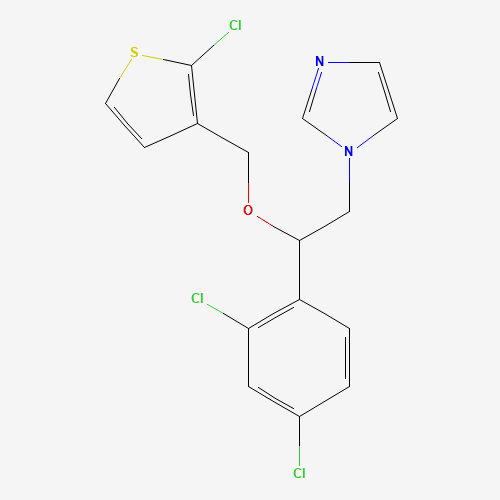 | Antifungal drug with antibacterial and anti-influenza activity (Clissold and Heel 1986; An et al. 2014) |
| 6 | DB00557 | Hydroxyzine | 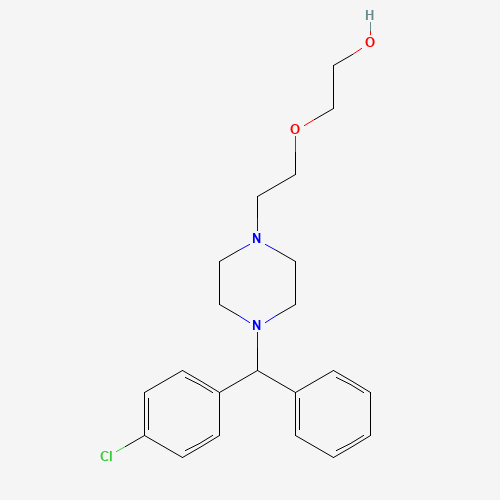 | Antihistamine drug with antibacterial and anti-influenza activity (An et al. 2014; Aybey et al. 2014) |
| 7 | DB01114 | Chlorpheniramine | 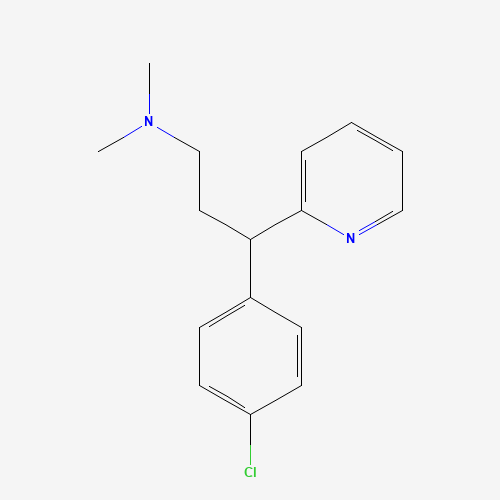 | Antihistamine drug with anti-influenza activity (Xu et al. 2018) |
| 8 | DB01624 | Zuclopenthixol | 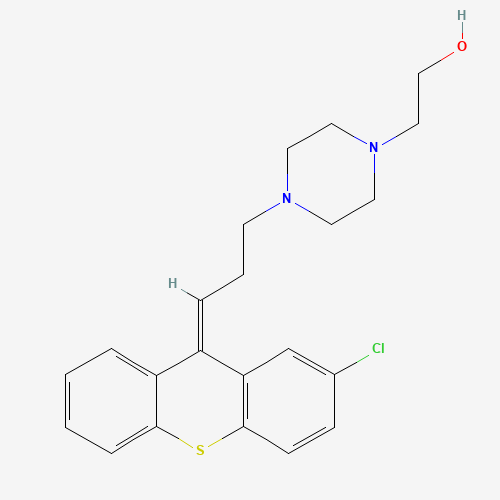 | Antipsychotic drug with antiviral activity (Ulferts et al. 2016) |
| 9 | DB01551 | Dihydrocodeine | 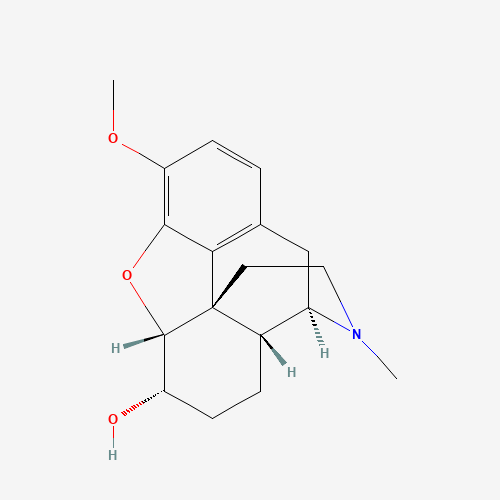 | Opioid analgesic prescribed in case of bacterial infections (Baker 2012) |
| 10 | DB08936 | Chlorcyclizine | 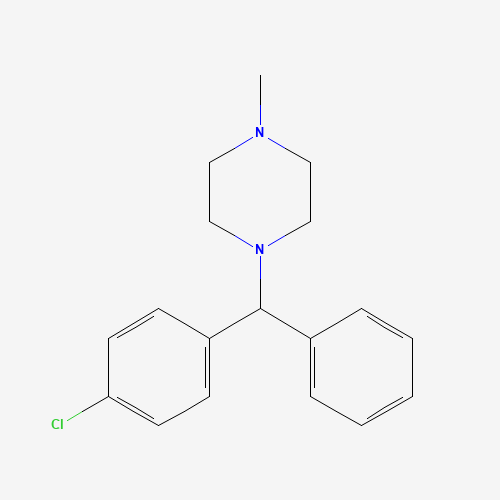 | Antihistamine drug and inhibitor of viral fusion in HCV (Hu et al. 2020) |

**Supplementary table S2** Two-dimensional structures of 10 top-scoring molecules docked with NA

| **SL. No** | **Drugbank ID** | **Molecule name** | **Structure** | **Current drug function** |
| --- | --- | --- | --- | --- |
| 1 | DB00284 | Acarbose | 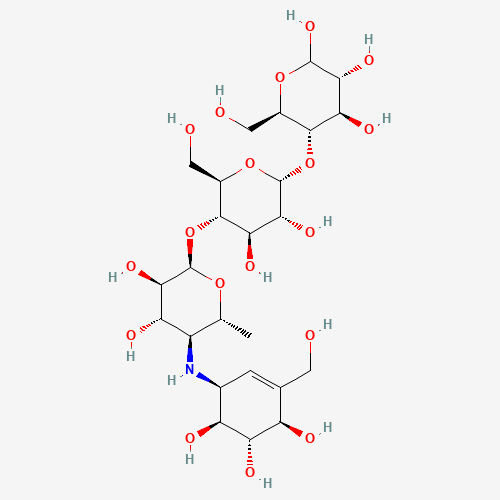 | An alpha-glucosidase inhibitor showing *in-vitro* and *in-vivo* activity against Enterovirus 71 and *in-silico* activity against SARS-CoV-2 (Kumar et al. 2020; Feng et al. 2020b; Sundar et al. 2021) |
| 2 | DB01698 | Rutin | 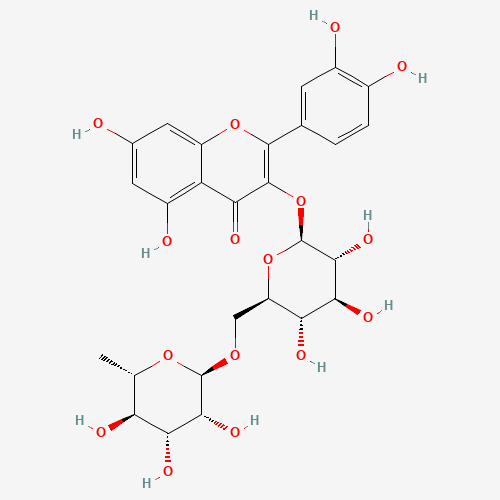 | Flavanoid with anti-influenza activity (Ling et al. 2020) |
| 3 | DB01421 | Paromomycin | 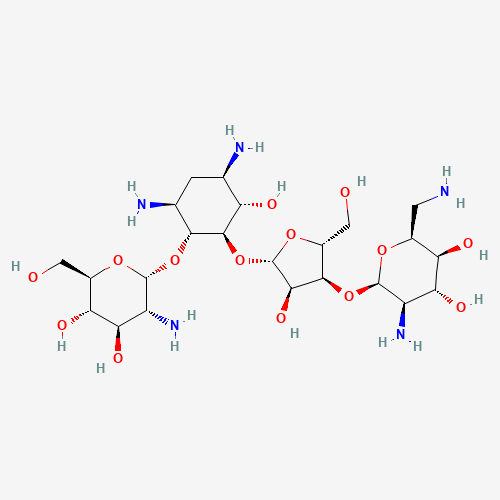 | Aminoglycoside antibiotic with anti-influenza activity (Kim et al. 2012) |
| 4 | DB01177 | Idarubicin | 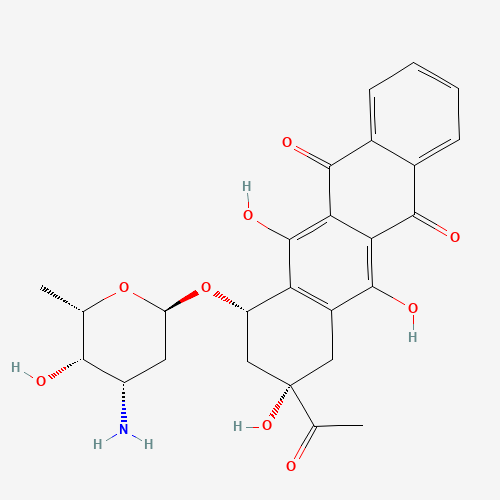 | Anticancer drug with antibacterial and antifungal activity (Kinnunen et al. 2009) |
| 5 | DB14726 | Dabigatran | 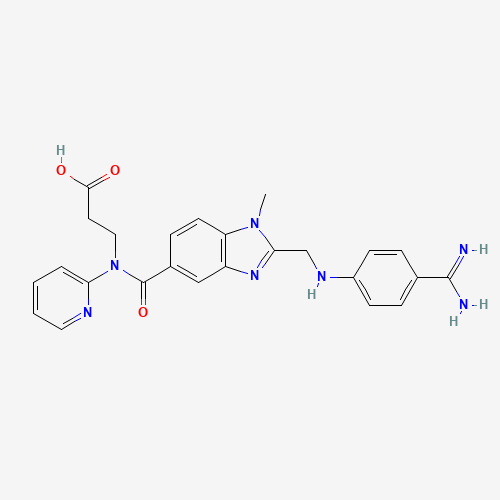 | Anticoagulant drug with antibacterial activity (Vanassche et al. 2011) |
| 6 | DB00198 | Oseltamivir | 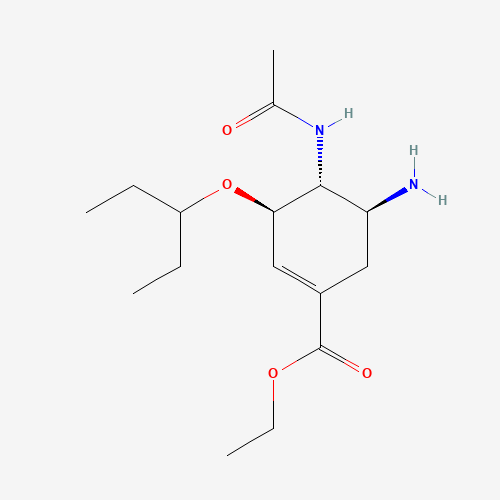 | Neuraminidase inhibitor in influenza virus (Laborda et al. 2016) |
| 7 | DB00140 | Riboflavin | 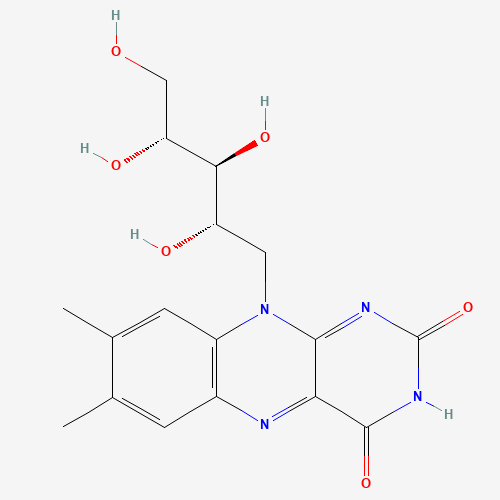 | A nutraceutical used for vitamin B2 deficiency with antibacterial and anti-viral activity (Ruane et al. 2004) |
| 8 | DB01136 | Carvedilol | 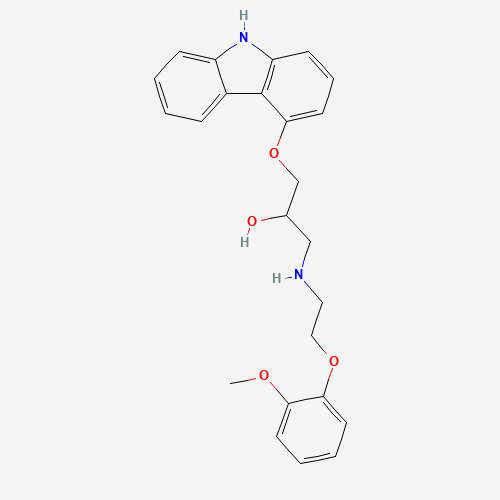 | An antihypertensive agent with antibacterial activity (Zawadzka et al. 2018) |
| 9 | DB00983 | Formoterol | 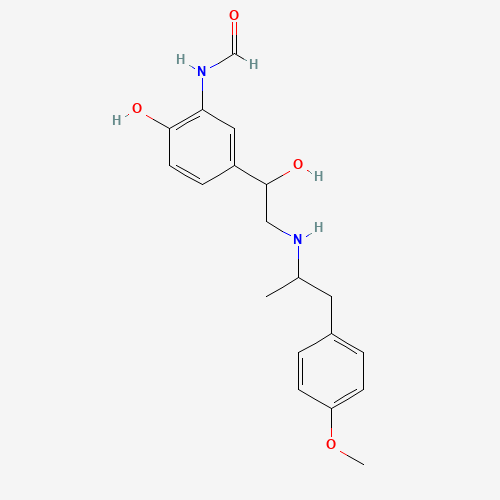 | Agonist of Beta-adrenergic receptor and enhances host defense against bacterial infections (Gross et al. 2009) |
| 10 | DB01259 | Lapatinib | 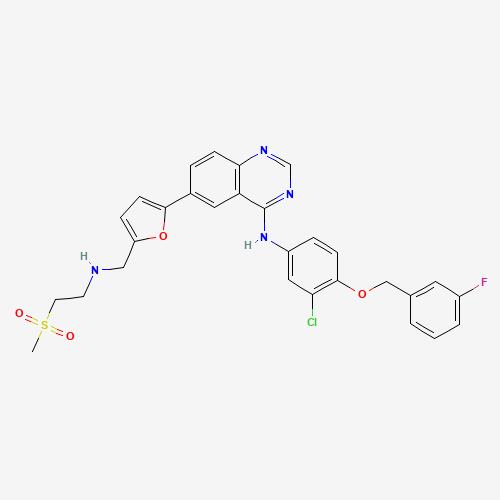 | Anticancer drug with antibacterial and antiviral activity (Raymonda et al. 2022; Liu et al. 2022) |
| Co-crystal ligand of 3TI6 (oseltamivir carboxylate) | | | 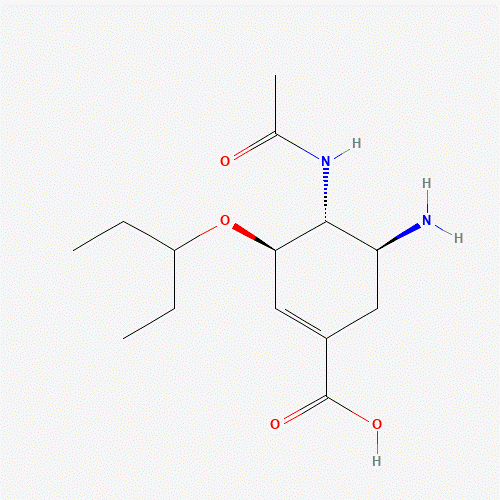 |  |
